# Selection analysis identifies unusual clustered mutational changes in Omicron lineage BA.1 that likely impact Spike function

**DOI:** 10.1101/2022.01.14.476382

**Authors:** Darren P Martin, Spyros Lytras, Alexander G Lucaci, Wolfgang Maier, Björn Grüning, Stephen D Shank, Steven Weaver, Oscar A MacLean, Richard J Orton, Philippe Lemey, Maciej F Boni, Houriiyah Tegally, Gordon Harkins, Cathrine Scheepers, Jinal N Bhiman, Josie Everatt, Daniel G Amoako, James Emmanuel San, Jennifer Giandhari, Alex Sigal, NGS-SA, Carolyn Williamson, Nei-yuan Hsiao, Anne von Gottberg, Arne De Klerk, Robert W Shafer, David L Robertson, Robert J Wilkinson, B Trevor Sewell, Richard Lessells, Anton Nekrutenko, Allison J. Greaney, Tyler N. Starr, Jesse D. Bloom, Ben Murrell, Eduan Wilkinson, Ravindra K Gupta, Tulio de Oliveira, Sergei L Kosakovsky Pond

## Abstract

Among the 30 non-synonymous nucleotide substitutions in the Omicron S-gene are 13 that have only rarely been seen in other SARS-CoV-2 sequences. These mutations cluster within three functionally important regions of the S-gene at sites that will likely impact (i) interactions between subunits of the Spike trimer and the predisposition of subunits to shift from down to up configurations, (ii) interactions of Spike with ACE2 receptors, and (iii) the priming of Spike for membrane fusion. We show here that, based on both the rarity of these 13 mutations in intrapatient sequencing reads and patterns of selection at the codon sites where the mutations occur in SARS-CoV-2 and related sarbecoviruses, prior to the emergence of Omicron the mutations would have been predicted to decrease the fitness of any genomes within which they occurred. We further propose that the mutations in each of the three clusters therefore cooperatively interact to both mitigate their individual fitness costs, and adaptively alter the function of Spike. Given the evident epidemic growth advantages of Omicron over all previously known SARS-CoV-2 lineages, it is crucial to determine both how such complex and highly adaptive mutation constellations were assembled within the Omicron S-gene, and why, despite unprecedented global genomic surveillance efforts, the early stages of this assembly process went completely undetected.

## Introduction

The Omicron (B.1.1.529) SARS-CoV-2 variant of concern (VOC) identified in Southern Africa in late November 2021 ^1^ is the product of extensive evolution within an infection context that has so far yielded at least three genetically distinct viral lineages (BA.1, BA.2 and BA.3) since it diverged from an ancestral B.1.1 lineage (presumably at some time in mid to late 2020). Three possible explanations for the sudden appearance of Omicron without any prior detection of intermediate/progenitor forms before its discovery are: (1) SARS-CoV-2 genomic surveillance in the region where Omicron originated might have been inadequate to detect intermediate forms; (2) long-term evolution in one or more chronically infected people - similar to the proposed origin of lineages such as Alpha and C.1.2 ^2 3 4^ - may have left intermediate forms unsampled within one or a few individual(s); and (3) reverse zoonosis to a non-human host, followed by undetected spread and diversification therein prior to spillover of some sub-lineages back into humans ^5^. At present there is no strong evidence to support or reject any of these hypotheses on the origin of Omicron, but as new data are collected, its origin may be more precisely identified.

Regardless of the route that Omicron took to eventual community transmission, the genome of the BA.1 lineage that caused surges of infections globally in late 2021 and early 2022, accumulated 53 mutations relative to the Wuhan-Hu-1 reference strain, with 30 non-synonymous substitutions in the Spike-encoding S-gene alone (Figure 1). Here, we characterize the selective pressures that may have acted during the genesis of the BA.1 lineage and curate available data on the likely adaptive value of the BA.1 S-gene mutations. We were particularly interested in identifying BA.1 S-gene codon sites displaying evolutionary patterns that differed from those of other SARS-CoV-2 lineages (including variation of SARS-CoV-2 in individual hosts), and closely related non-human sarbecoviruses. We use these comparisons to identify which BA.1 S-gene mutations might contribute to recently discovered shifts relative to other SARS-CoV-2 variants in the way that BA.1 interacts with human and animal ACE2 receptors and is primed by cellular proteases to mediate cellular entry ^6–10^. Our analysis identifies three clustered sets of mutations in the Spike protein, involving amino acids substitutions at 13 sites previously highly conserved across other SARS-CoV-2 lineages and other sarbecoviruses. The dramatic about-face in evolutionary dynamics at the 13 codon sites encoding these amino acids indicates that the mutations at these sites in BA.1 are likely interacting with one another, that the combined effects of these interactions are likely adaptive, and that these adaptations likely underlie at least some of the recently discovered shifts in BA.1 Spike function.

**Figure 1.**
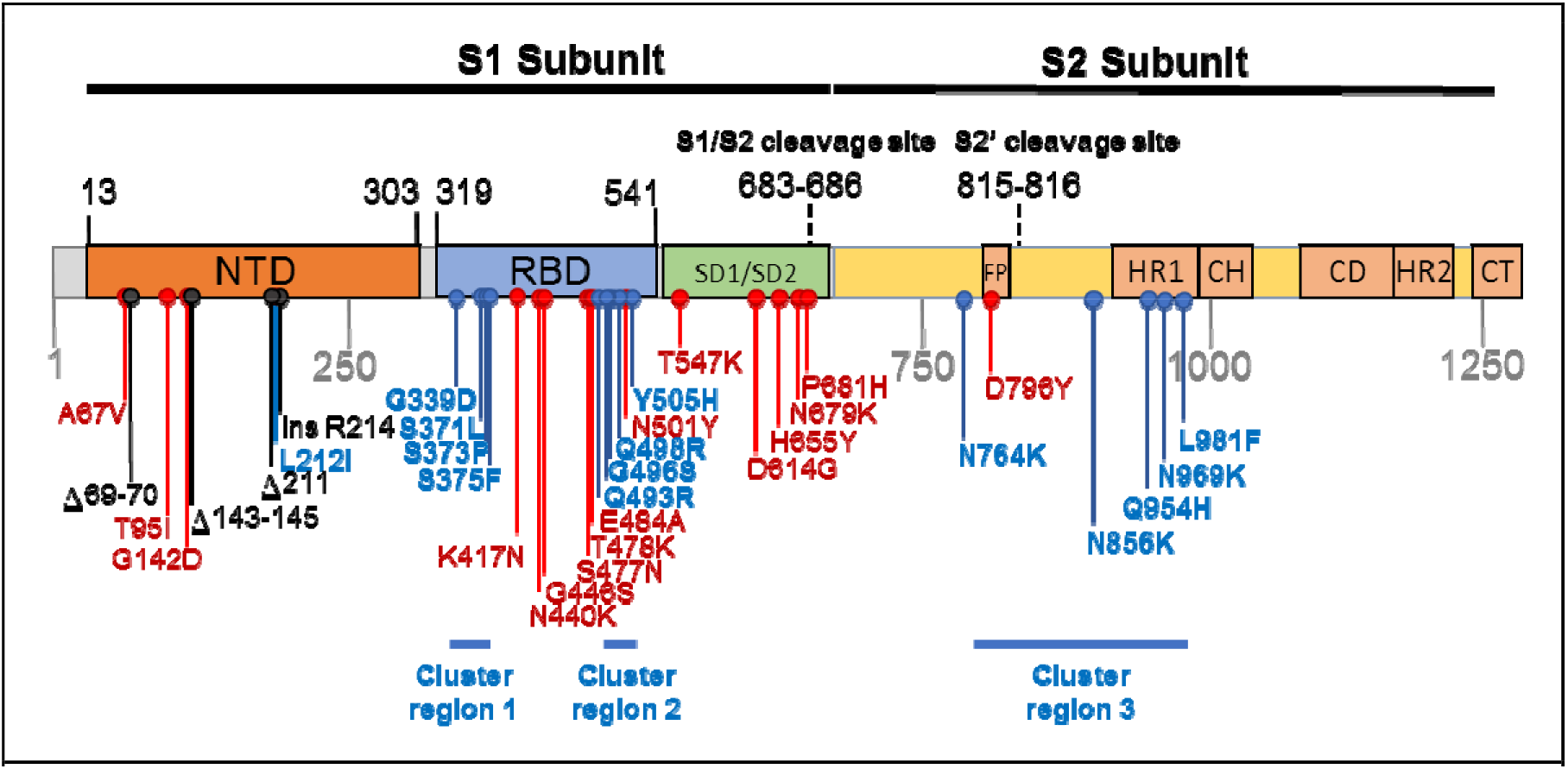
Mutations characterising the S-gene of the BA.1 lineage viruses. Amino acid changes resulting from non-synonymous substitutions relative to the Wuhan-Hu-1 sequence are indicated in: Blue = those attributable to nucleotide substitutions at codon sites that are either negatively selected or are evolving under no detectable selection in non-Omicron sequences and cluster within three regions labelled here as cluster regions 1, 2 and 3; Red = those attributable to nucleotide substitutions at codon sites that are detectably evolving under positive selection in non-Omicron sequences; and Black = those attributable to insertion and deletion mutations. NTD = N-terminal domain; RBD = receptor binding domain; SD1/SD2 = subdomain 1 and 2; FP= fusion peptide, HR1 = heptad repeat 1; CH =central helix; CD = connector domain; HR2 = heptad repeat 2; CT = cytoplasmic tail.

## Results and Discussion

### Many of the BA.1 S-gene mutations likely contribute to viral adaptation

Relative to the Wuhan-Hu-1 reference variant of SARS-CoV-2, BA.1 has 30 non-synonymous substitutions in its S-gene (Figure 1). Sixteen of the codon sites where these mutations occur are presently, or have recently been, detectably evolving under positive selection when considering all SARS-CoV-2 genomic data prior to the discovery of Omicron (Table 1, Figure 2, https://observablehq.com/@spond/selection-profile). For context, this fraction of positively selected sites (0.53) is approximately four times higher than the fraction of all SARS-CoV-2 S-gene sites that have ever shown any signals of positive selection (0.14).

**Table 1.**
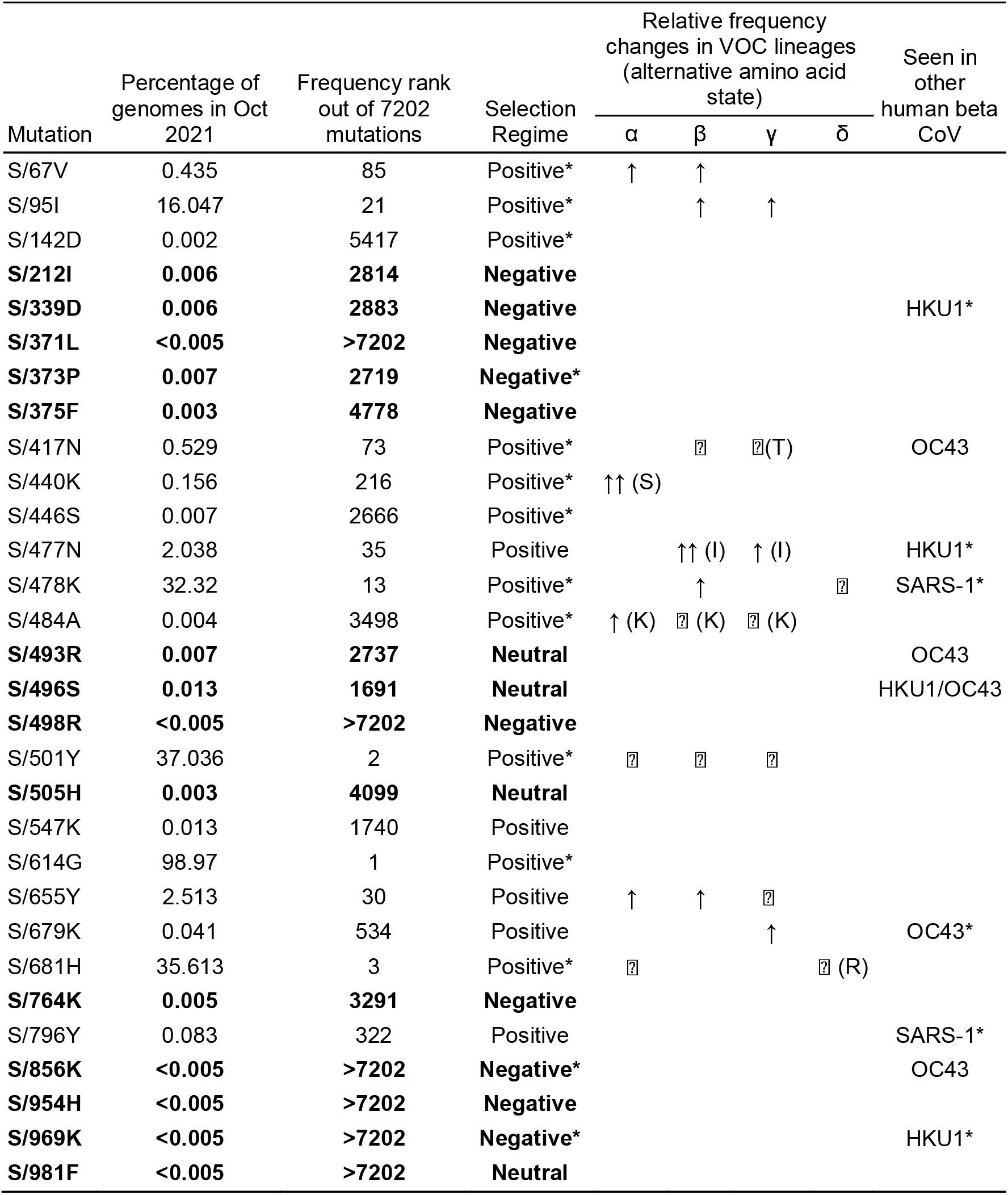
Frequencies in non-Omicron SARS-CoV-2 genomes of non-synonymous mutations seen in the S-gene of BA.1. Rows in bold indicate mutations at previously negatively selected or neutrally evolving sites. VOC columns track fold changes in mutation frequencies at corresponding sites in other VOCs (before and after boundaries are defined to create somewhat balanced sizes of sequence sets; the boundary is 2021/04/15 for α,β,γ and 2021/06/01 for δ). If another amino-acid residue is included in parentheses, then this residue has increased in frequency at the same site. ↑ - 2-10x fold increase ↑↑ - >10x fold increase. ⍰ - lineage defining/majority mutation. (*) in other human beta-CoV - consensus residue in species matches the BA.1 residue; based on the sequence alignment from ^11^

| Mutation | Percentage of genomes in Oct 2021 | Frequency rank out of 7202 mutations | Selection Regime | Relative frequency changes in VOC lineages (alternative amino acid state) |  |  |  | Seen in other human beta CoV |
| --- | --- | --- | --- | --- | --- | --- | --- | --- |
| | | | | $\alpha$ | $\beta$ | $\gamma$ | $\delta$ | |
| S/67V | 0.435 | 85 | Positive* | ↑ | ↑ | | | |
| S/95I | 16.047 | 21 | Positive* | | ↑ | ↑ | | |
| S/142D | 0.002 | 5417 | Positive* |  |  |  |  |  |
| **S/212I** | **0.006** | **2814** | **Negative** |  |  |  |  |  |
| **S/339D** | **0.006** | **2883** | **Negative** |  |  |  |  | HKU1* |
| **S/371L** | **<0.005** | **>7202** | **Negative** |  |  |  |  |  |
| **S/373P** | **0.007** | **2719** | **Negative\*** |  |  |  |  |  |
| **S/375F** | **0.003** | **4778** | **Negative** |  |  |  |  |  |
| S/417N | 0.529 | 73 | Positive* | | □ | □(T) | | OC43 |
| S/440K | 0.156 | 216 | Positive* | $\uparrow\uparrow$ (S) | | | | |
| S/446S | 0.007 | 2666 | Positive* |  |  |  |  |  |
| S/477N | 2.038 | 35 | Positive | | $\uparrow\uparrow$ (I) | ↑ (I) | | HKU1* |
| S/478K | 32.32 | 13 | Positive* | | ↑ | | □ | SARS-1* |
| S/484A | 0.004 | 3498 | Positive* | ↑ (K) | □ (K) | □ (K) | | |
| **S/493R** | **0.007** | **2737** | **Neutral** |  |  |  |  | OC43 |
| **S/496S** | **0.013** | **1691** | **Neutral** |  |  |  |  | HKU1/OC43 |
| **S/498R** | **<0.005** | **>7202** | **Negative** |  |  |  |  |  |
| S/501Y | 37.036 | 2 | Positive* | □ | □ | □ | | |
| **S/505H** | **0.003** | **4099** | **Neutral** |  |  |  |  |  |
| S/547K | 0.013 | 1740 | Positive |  |  |  |  |  |
| S/614G | 98.97 | 1 | Positive* |  |  |  |  |  |
| S/655Y | 2.513 | 30 | Positive | ↑ | ↑ | □ | | |
| S/679K | 0.041 | 534 | Positive | | | ↑ | | OC43* |
| S/681H | 35.613 | 3 | Positive* | □ | | | □ (R) | |
| **S/764K** | **0.005** | **3291** | **Negative** |  |  |  |  |  |
| S/796Y | 0.083 | 322 | Positive |  |  |  |  | SARS-1* |
| **S/856K** | **<0.005** | **>7202** | **Negative\*** |  |  |  |  | OC43 |
| **S/954H** | **<0.005** | **>7202** | **Negative** |  |  |  |  |  |
| **S/969K** | **<0.005** | **>7202** | **Negative\*** |  |  |  |  | HKU1* |
| **S/981F** | **<0.005** | **>7202** | **Neutral** |  |  |  |  |  |

The observed substitutions at four of these sixteen sites (K417N ^16^, N501Y ^17–19^, H655Y, P681H ^20^) and a two-nucleotide deletion at one additional site (Δ69-70 ^21^) are among the nineteen “501Y meta-signature” Spike mutations that are likely highly adaptive within the context of 501Y lineage viruses such as the Alpha, Beta and Gamma VOCs ^15^. Given that the BA.1 mutations at these sites converge on those seen in these other VOCs, they are likely to be adaptive in BA.1 lineage viruses as well (sites coloured red in Figure 3).

**Figure 2.**
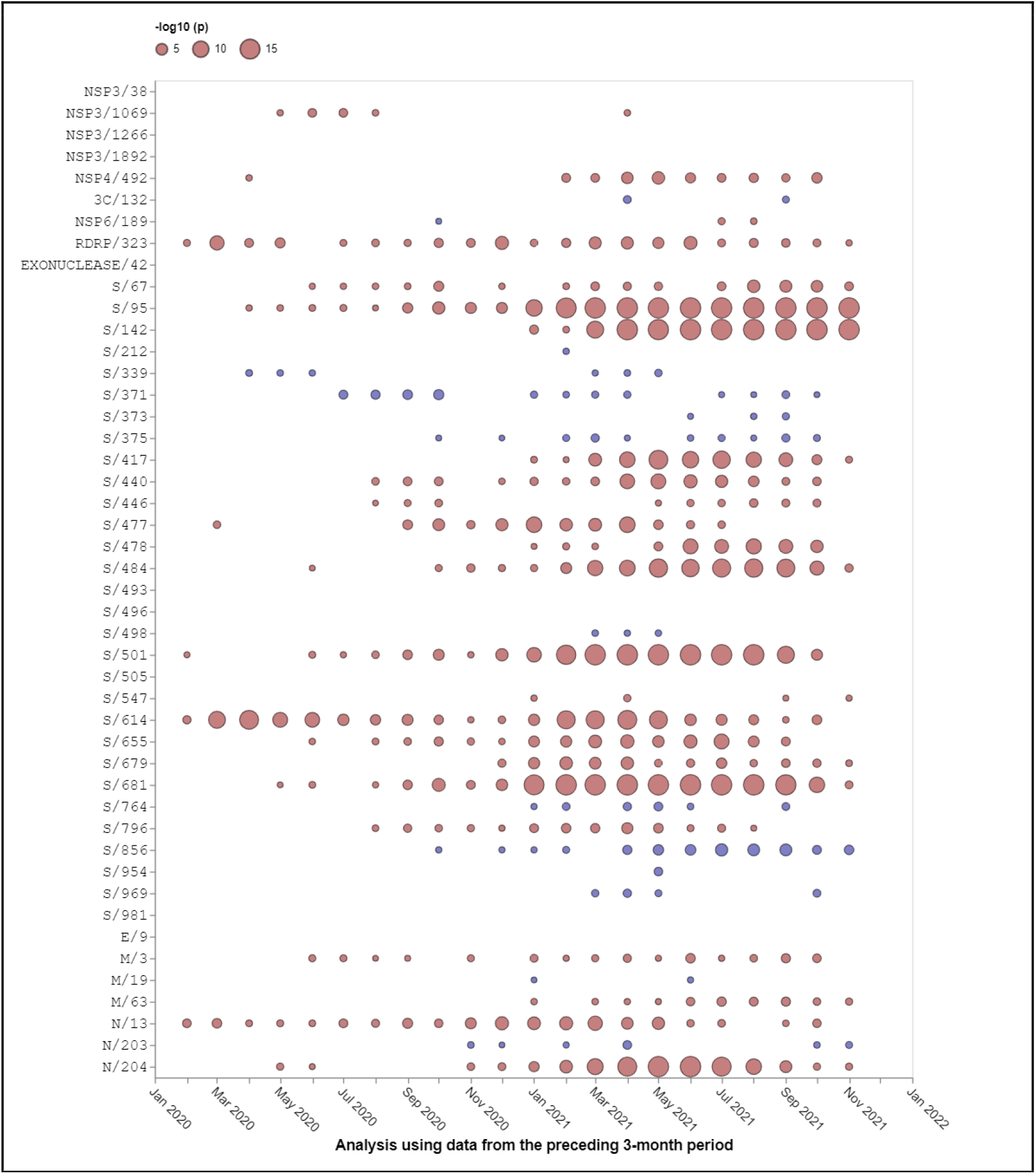
Selection signals that were evident at BA.1 amino acid change sites in other SARS-CoV-2 lineages prior to the emergence of Omicron. All SARS-CoV-2 near full-length genome sequences present in GISAID ^12^ on 21 November 2021 that passed various quality control checks were split up into three month sampling windows and analysed using the FEL method restricted to internal tree branches ^13^ implemented in Hyphy 2.5 ^14^. This method was also used in ^15^. Red circles show sites under positive selection (selection favouring changes at amino acid states encoded at these sites). Blue circles show sites under negative selection (selection against non-synonymous changes). When no circle is shown, the corresponding site offered no statistical evidence for nonneutral evolution at a given time point. The areas of circles indicate the statistical strength of the selection signal (and not the actual strength of selection) within sequences sampled in the three months preceding the 1st day of the indicated months. Note that none of these analyses included any Omicron sequences, hence selection signals are derived solely from other SARS-CoV-2 lineages.

**Figure 3.**
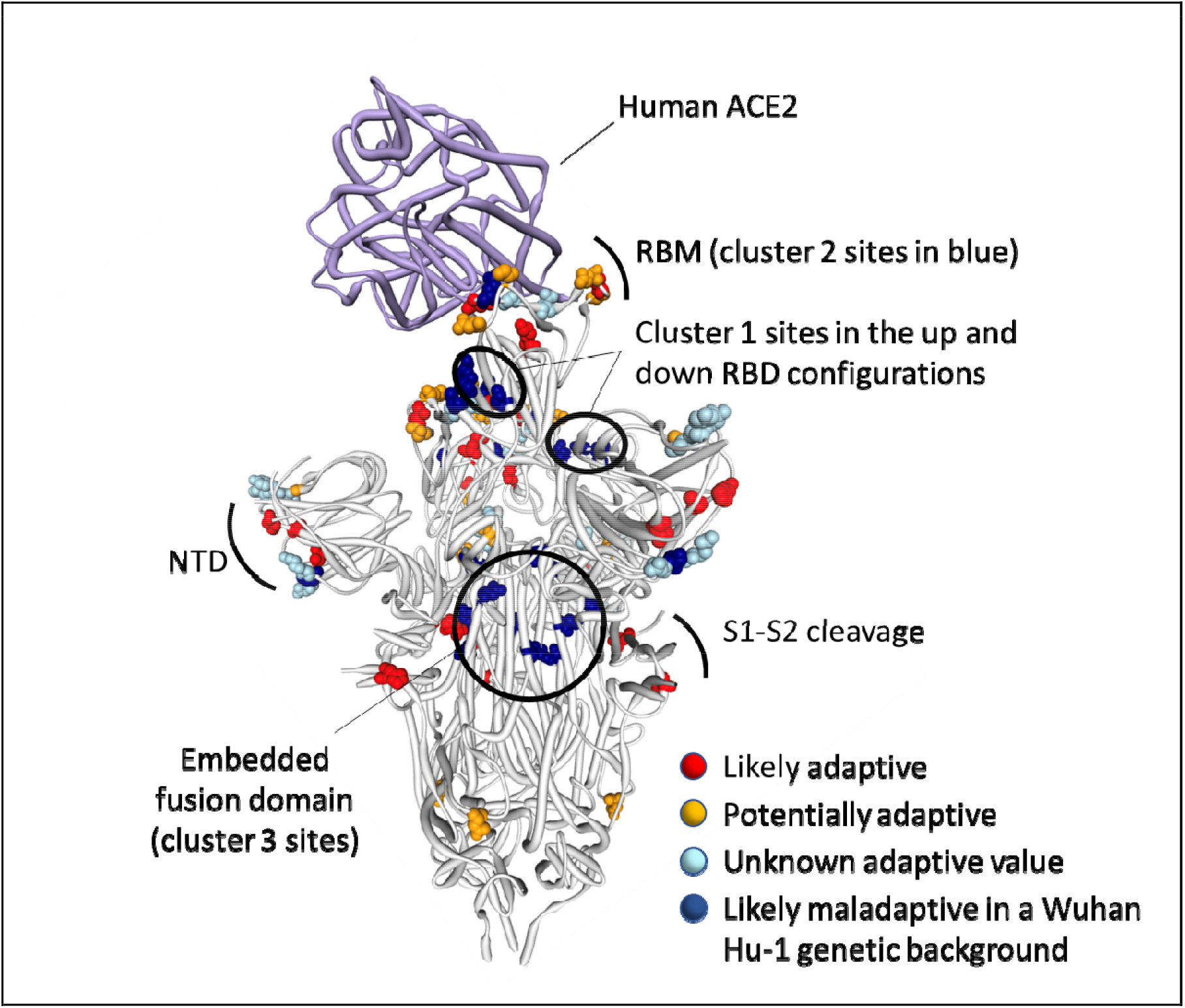
Distribution of BA.1 amino acid replacements on the three dimensional SARS-CoV-2 Spike trimer. In this rendering of the trimer, one subunit is shown in the “up” or “open” configuration while interacting with human ACE2 ^23^. The other two subunits are in the “down” or “closed” configurations. Amino acids are color coded according to their likely contribution to viral adaptation in a Wuhan-Hu-1-like genetic background based on (1) patterns of synonymous and nonsynonymous substitutions at the codons encoding these amino acids in non-Omicron sequences, (2) patterns of mutational convergence between viruses in different VOCs and (3) increases in the frequency over time of VOC sub-lineages encoding amino acids that match those found in BA.1. NTD = N-terminal domain, RBD = Receptor binding domain; RBM = receptor binding motif. Locations of sites in the three clusters of BA.1 mutations that are rarely seen and fall at either negatively selected (dark blue) and neutrally evolving (light blue) sites. An interactive version of this figure can be found here: https://observablehq.com/@stephenshank/sars-cov-2-ace2-protein-interaction-and-evolution-for-omicr

A further four BA.1 S-gene mutations are found in SARS-CoV-2 sequences belonging to other VOC lineages, and are either VOC lineage defining mutations (majority mutations), or are lower frequency mutations that have increased in frequency >2 fold between early and late VOC lineage circulation periods within sampled sequences belonging to these lineages (A67V in Alpha and Beta, T95I in Beta and Gamma, T478K in Beta, and N679K in Gamma; https://observablehq.com/@spond/sc2-selection-trends): an indication that these mutations too are likely adaptive in BA.1 lineage viruses (Table 1). Additionally, three other BA.1 S-gene mutations either: (1) occur at the same codon sites as Alpha, Beta, Gamma or Delta lineage defining mutations but encode a different amino acid than these other lineages (E484A in BA.1 and E484K in Beta and Gamma); or (2) occur at the same codon sites as mutations in VOC lineages that increased in frequency > 2 fold between early and late VOC lineage circulation periods but encode a different amino acid than these other lineages (N440K in BA.1 and N440S in Alpha; S477N in BA.1 and S477I in Beta and Gamma). Lastly, the S/D796Y mutation occurs at one of the four sites identified as potential locations of adaptation in human beta-coronaviruses via the analysis of convergent evolutionary patterns and functional impact (Table 1)^11^ and a mutation at this site has previously been inferred to be potentially adaptive within the context of a chronic SARS-CoV-2 infection ^22^. All of these mutations likely have a substantial impact on the phenotype of BA.1 lineage viruses (coloured orange in Figure 3).

Finally, three deletions (Δ69-70, Δ143-145 and Δ211-212) and a nine nucleotide insertion (between codons 214 and 215) in the N-terminal domain encoding part of the S-gene all likely have phenotypic impacts and all are potentially adaptive but are not considered further here because they are not amenable to analysis by natural selection analysis methods that focus on patterns of synonymous and non-synonymous mutations.

### Clusters of BA.1 mutations occur at neutral or negatively selected S-gene sites

The mutations occurring at the 14 BA.1 Spike codons which display either evidence of negative selection or no evidence of selection (neutral evolution), have rarely been seen within previously sampled sequences (bold rows in Table 1; https://observablehq.com/@spond/omicron-mutations-tables) indicating the action of strong purifying selection due to functional constraints. Despite the rarity of these mutations in assembled genomes, it is not uncommon to find them in within-patient sequence datasets (Figure 4), often at sub-consensus allelic frequencies. This indicates that, with the possible exceptions of S/S371L, S/N764K, S/N856K and S/Q954H, the mutations at these sites are not rare simply because they are unlikely to occur (note the sizes and numbers of dots corresponding to these mutations in Figure 4), but rather because whenever they do occur they are unlikely to either increase sufficiently in frequency to be transmitted (note the predominantly light orange/yellow colours of the dots corresponding to these mutations in Figure 4), or increase sufficiently in frequency among transmitting viruses to be detected by genomic surveillance.

**Figure 4.**
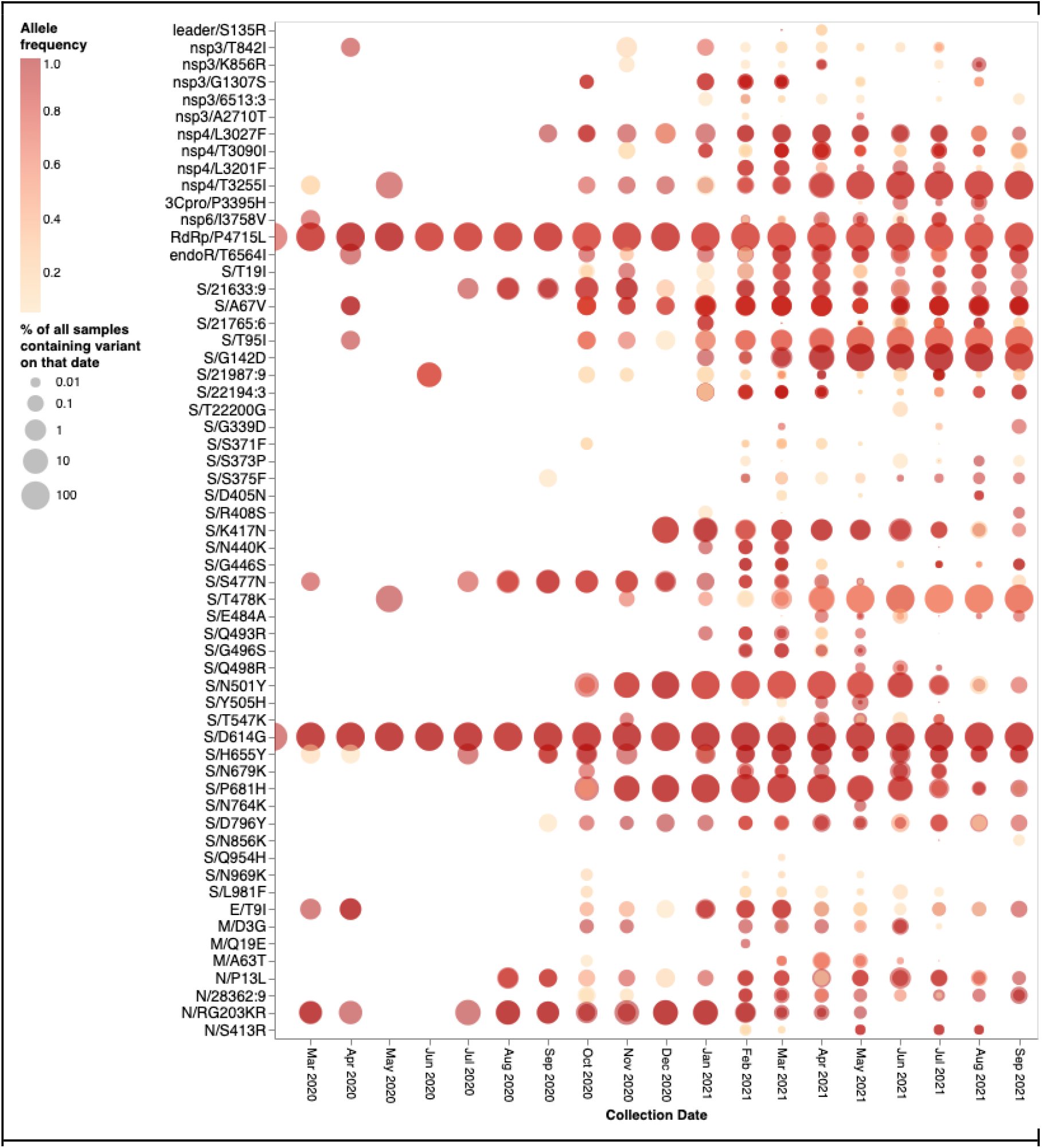
Intrapatient allelic variation seen at BA.1 amino acid mutation sites in a subset of SARS-CoV-2 raw sequencing data since March 2020 analyzed using a standardized variant calling pipeline ^24^. The areas of the circles indicate the proportions of raw sequence datasets (per 1,000 samples) where a mutation away from the Wuhan-Hu-1 consensus sequence was called. The colour of the circle indicates the median intrapatient allele frequency (AF) in datasets for which each mutation was detected. Mutations occurring at lower AFs are only present in a subpopulation of viruses in a particular host. The data has been generated by calling variants from read-level data of 230,506 samples from COG-UK, Estonia, Greece, Ireland, and South Africa: PRJEB37886, PRJEB42961 (and multiple other bioprojects with the study title: Whole genome sequencing of SARS-CoV-2 from Covid-19 patients from Estonia), PRJEB44141, PRJEB40277 and PRJNA636748. Note that S371L is the result of two nucleotide substitutions in codon S/371 and was never detected in intrapatient samples. S371F represents an intermediate mutation between the Wuhan-Hu-1 state and that of BA.1 and is presented here for completeness.

On their own, none of these 14 BA.1 mutations at codon sites that have previously been evolving either neutrally or under negative selection prior to November 2021 would be expected to provide SARS-CoV-2 with any selective advantage. If the BA.1 mutations observed at the ten negatively selected S-gene codon sites had occurred in the Wuhan-Hu-1 sequence, it is very likely that they would have been selected against. Specifically, since the start of the pandemic Spike proteins tended to function best whenever they had amino acids at these ten sites that were the same as those in the Spike encoded by the Wuhan-Hu-1 sequence.

It is clear that the amino acids encoded by 13 of the 14 mutated codon sites in the BA.1 S-gene that either show evidence of negative selection or no evidence of any selection, cluster within three regions of the Spike three dimensional structure (dark blue sites in Figure 3):

1. **Cluster region 1** in the RBD (green sites in Figure 5): codons/amino acids S/339, S/371, S/373 and S/375; may be targeted by some class 4 neutralizing antibodies ^25^. S/371L alone impacts, but probably does not provide escape from, binding of some antibodies in all four neutralizing antibody classes ^26^ suggesting that, in a Wuhan-Hu-1 genetic background, it may substantially impact the trimerization, glycosylation profile, or balance of up-down conformations of Spike.
2. **Cluster region 2** in the RBM (cyan sites in Figure 5) including codons/amino acids S/493, S/496, S/498, and S/505. This region is known to be targeted by class 1 and class 2 neutralizing antibodies ^16,27^. S/493 is, in fact, a known target of such antibodies. Accordingly S/Q493R (as occurs in BA.1), escapes some class 2 neutralizing antibodies ^26^, S/Q493R and S/Q493K escape mutations have been selected in VSV *in vitro* experiments^28^, and the S/Q493K mutation has arisen previously in the context of a chronic SARS-CoV-2 infection^29^. The S/Q498R and S/Q493R mutations also yield two additional salt bridges when binding human ACE2 ^10,30^ and it is likely that the increased affinity of BA.1 Spike for human ACE2 relative to that of Alpha, Beta, Delta and Wuhan-Hu-1 ^7–9^ will further decrease its sensitivity to neutralisation.
3. **Cluster region 3** in the fusion domain (yellow sites in Figure 5): codons/amino acids S/764, S/856, S/954, S/969, S/981; a region of Spike currently not known to be targeted by neutralizing antibodies. The S/N764K, S/N856K and S/N969K mutations are likely to enhance interactions between the S1 and S2 subunits of the BA.1 Spike and are likely to contribute to reduced S1 shedding following proteolytic cleavage of the polybasic S1/S2 site ^10,31^

**Figure 5.**
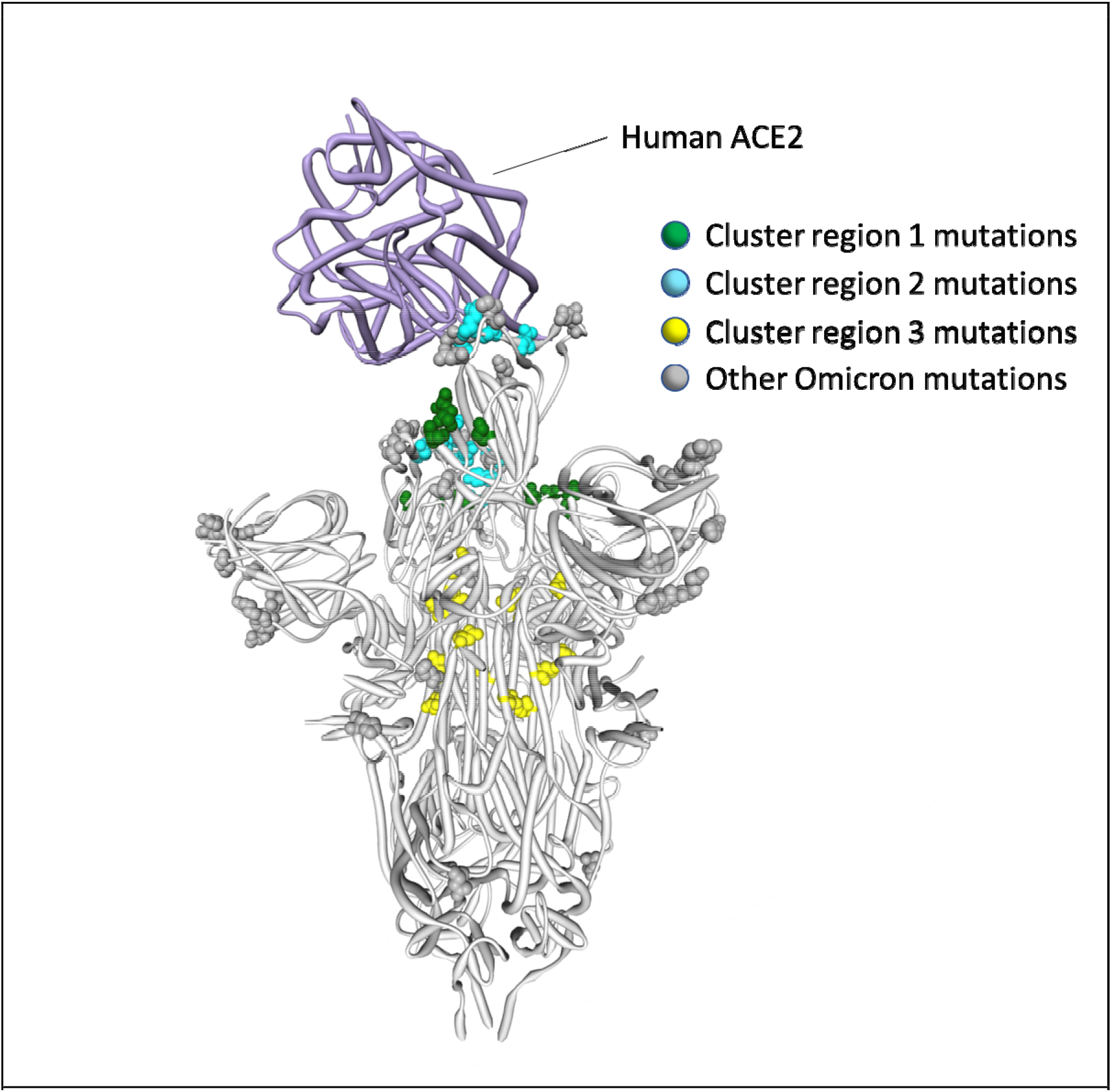
Positions on the three dimensional SARS-CoV-2 Spike trimer of amino acids encoded by three clusters of BA.1 codon sites that are evolving either neutrally or under negative selection in non-Omicron SARS-CoV-2 sequences. The Spike subunit interacting with human ACE2 is in the “up” configuration and the other two are in the “down” configuration ^23^. The cluster region 1 and 2 encoded amino acid changes in BA.1 (in green and blue respectively) are within the receptor binding domain of Spike with the cluster 2 encoded changes located within the receptor binding motif. The cluster region 3 mutations are within the fusion domain of Spike. An interactive version of this figure can be found at https://observablehq.com/@stephenshank/sc2-omicron-clusters.

### Selection patterns in sarbecoviruses confirm that, on their own, many BA.1 mutations would likely be deleterious

To determine whether patterns of selection at the Omicron/BA.1-specific sites are broadly consistent with those occurring in the horseshoe bat-infecting SARS-related coronaviruses, in the *Sarbecovirus* subgenus to which SARS-CoV-2 belongs, we examined patterns of synonymous and non-synonymous substitutions in 167 publicly available Sarbecovirus genomes. Accounting for recombination, we tested for selection signatures at all 44 codons encoding amino acids that differ between Wuhan-Hu-1 and BA.1 (https://observablehq.com/@spond/ncos-evolution-nov-2021). We specifically focused the analyses on selection signals in the subset of sarbecoviruses that are more closely related to SARS-CoV-2 in each recombination-free part of their genome: a group of sequences we refer to as the nCoV clade ^32^. Depending on the recombination-free genome region being considered, this clade was represented by between 15 and 27 sequences. We refer to the remaining sarbecoviruses as the non-nCoV sequences.

Of the 44 codon sites considered, 26 are detectably evolving under negative selection (FEL p-value <0.05; ^14 13^) and one (S/417) under positive selection (MEME p-value <0.05; ^33^) in the nCoV clade. This positive selection signal at S/417 reflects an encoded amino acid change from an ancestral V that is present in all background sequences, to a K that is specific to the nCoV clade. A K is also encoded at this site in Wuhan-Hu-1 but has since changed multiple times in various SARS-CoV-2 lineages: for example, to an N during the genesis of lineages such as Omicron and Beta and to a T during the genesis of the Gamma lineage.

We were, however, particularly interested in whether the cluster 1, 2 and 3 mutation sites in the S-gene were also evolving in a constrained manner (i.e., under negative selection) in the nCoV clade and, if so, what the selectively favoured encoded amino acid states were at these sites. Consistent with the hypothesis that the Wuhan-Hu-1 encoded amino acid states are generally constrained in the closest known SARS-CoV-2 relatives, the cluster 1 sites S/339, S/373 and S/375, the cluster 2 site S/505 and the cluster 3 sites S/764, S/856, S/969 and S/981 were all detectably evolving under negative selection in the nCoV clade viruses with the Wuhan-Hu-1 encoded amino acid state being favoured at all eight of the sites. Also consistent with the hypothesis, two of the remaining five sites across the clusters that were not detectably evolving under negative selection in the nCoV clade (S/371 and S/954) predominantly encoded the Wuhan-Hu-1 amino acid state in all sarbecoviruses. Only the cluster 2 sites S/493, S/496 and S/498 seem to vary substantially across the *Sarbecovirus* subgenus.

### What can the sarbecoviruses tell us about the biological consequences of the rarely seen BA.1 mutations?

Despite the observation that, even among sarbecoviruses, BA.1 mutations seen in cluster regions 1, 2 and 3 are only rarely seen, the instances where they do occur might be illuminating. For example, among the bat-infecting sarbecoviruses, the BA.1 S/G339D substitution (in cluster region 1) has primarily to date been found among the bat-infecting viruses within a clade (Figure 6) that does not use ACE2 as a cell entry receptor ^34^. The change in receptor binding function in these viruses is, however, most likely due to two RBM deletions that are also specific to this clade. Further, cluster region 1 codon sites S/371, S/373 and S/375 encode a conserved serine (S) in almost all the analysed sarbecoviruses (164/167, 165/167 and 167/167 respectively). The change at sites S/371 and S/375 from an encoded polar residue (S) to a hydrophobic residue (an L at S/371 and an F at S/375) implies a substantial change in the biochemical properties of this region of Spike that has never before been seen in any sarbecovirus. These changes could be associated with SARS-CoV-2’s unique loss of N370 glycosylation site relative to all other sarbecoviruses ^35^, or packing of this surface with other BA.1 changes in cluster 2 (e.g. S/Y505H) in the locked Spike trimer structure.

**Figure 6.**
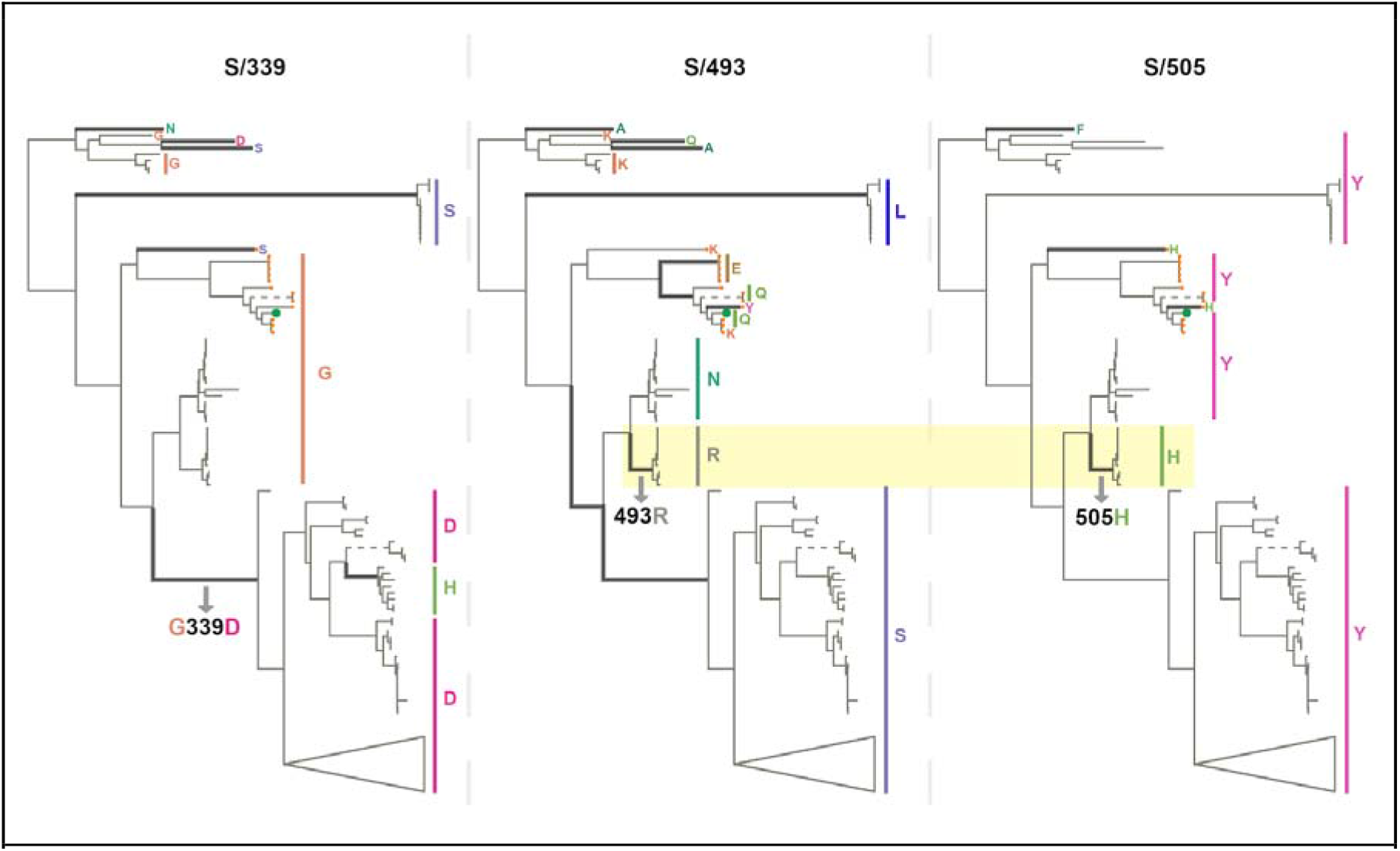
Phylogenetic trees of 167 sarbecoviruses indicating patterns of selection at S-gene codons S/339 (left tree), S/493 (middle tree) and S/505 (right tree). Branches along which amino acid states have changed are indicated with thick lines. Dashed lines represent long branches that have been shortened for visual clarity. The highlighted segments of the middle and right trees indicate the branch across which S/N493R and S/Y505H mutations occurred. The trees represent evolutionary relationships between putatively non-recombinant sequence fragments in the genome region corresponding to Wuhan-Hu-1 Spike positions 324-654. The clade containing sarbecoviruses sampled in Europe and Africa has been used as the outgroup for rooting. Tree tips are annotated by amino acid states at the respective sites. SARS-CoV-2 is annotated with a green tip symbol and the nCoV clade sequences with a tip symbol in orange.

As with SARS-CoV-2, the amino acids encoded at cluster region 2 sites (all of which fall within the RBM) vary substantially between different sarbecoviruses but without any associated signals of positive selection at these sites within the nCoV clade. Notably, the same BA.1 encoded amino acids at codon S/493R and S/505H also co-occur in a clade of sarbecoviruses that are closely related to SARS-CoV (virus accessions: KY417144, OK017858, KY417146, OK017852, OK017855, OK017853, OK017854, OK017856, OK017857); although S/493R (AY613951 and AY613948) and S/505H (MN996532, LC556375) can also occur independently. Besides the various Omicron sublineages, S/493R and S/505H are not found as a pair in any SARS-CoV-2 sequences. These mutations occurring along the same branch of the sarbecovirus tree (Figure 6) suggests that, rather than favouring changes at the sites individually, selection may favour simultaneous changes to S/493R and S/505H due to these residues together having a greater combined fitness benefit than the sum of their individual effects: a type of interaction between genome sites referred to as positive epistasis.

The region 3 cluster sites are conserved across the sarbecoviruses with almost all known viruses having the same residues at these sites as the Wuhan-Hu-1 SARS-CoV-2 strain. This supports the hypothesis that, when considered individually, the mutations seen at these fusion domain sites in BA.1 are likely to be maladaptive.

### BA.1 mutations at neutral or negatively selected S-gene sites might only be adaptive when they co-occur

Given both the apparent selective constraints on mutations arising at the cluster region 1, 2 and 3 sites in SARS-CoV-2 and other sarbecoviruses, and the rarity of observed mutations at these sites among the millions of assembled SARS-CoV-2 genomes (despite evidence that individually such mutations do regularly occur during within-host evolution; Figure 4), it is very likely that BA.1 mutations at cluster region, 1, 2 and 3 sites are maladaptive when present on their own. Nevertheless, the presence of mutations at these sites in BA.1, a lineage of viruses that is clearly highly adapted, suggests that these mutations might interact with one another such that, when present together, they become adaptive. Therefore while individually the mutations might decrease the fitness of any genome in which they occur, collectively they might compensate for one-another’s deficits to yield a fitter virus genotype.

Positive epistasis of this type has, in fact, already been demonstrated between the cluster 2 mutation, S/Q498R, and the pivotal mutation of the 501Y SARS-CoV-2 lineages, S/N501Y. Whereas S/498R only marginally impacts the affinity of Spike for human ACE2 when present with S/501N ^17^, it increases ACE2 binding affinity approximately four-fold when present with S/501Y ^19,36^.

Further, structural analyses of Spike trimer interactions imply that epistasis likely occurs among and between some cluster 1 and cluster 2 mutation sites. Specifically, in the context of Wuhan-Hu-1, S/371S and S/373S (among the cluster 1 mutation sites) of one Spike subunit within a trimer, are likely to interact via hydrogen bonds with S/493Q and S/505Y (among the cluster 2 mutation sites) of another subunit in the trimer when Spike is in its down configuration (Figure 5; ^37^). These interactions likely contribute to how readily Spike transitions to its up configuration and, in this regard, the S/S371L, S/S373P cluster 1 mutations and the S/Q493R and S/Y505H cluster 2 mutations in BA.1 may collectively destabilise the down configuration (such as occurs with the S/D614G mutation ^38^), to optimize ACE2 binding ^39^. If the S/S371L mutation on its own does indeed contribute to destabilizing the down configuration of Spike, then this might explain why, within the context of the Wuhan-Hu-1 sequence at least, this mutation results in moderate reduction in neutralization by monoclonal antibodies in all four RBD neutralizing antibody classes; including those that do not have S/371 in their binding footprint ^40^.

If mutations in the three cluster regions do epistatically interact with one another, then one might expect that selection would favour their co-occurrence either within individual SARS-CoV-2 genome sequences that have so far been sampled, or as minor variants within unassembled intrapatient sequence data. We failed to detect such associations in any systematic manner (Figure 7). While there are individual pairs of BA.1 mutations that cooccur more frequently than expected by chance (e.g. 440K in the presence of T95I), they do not involve cluster 1, 2, and 3 mutations. Furthermore, many of the BA.1 mutation pairs occur together **less** frequently than expected by chance (e.g. 478K and 501Y). Rather than reflecting an absence of epistasis between the cluster 1, 2, and 3 mutation sites our failure to detect the co-occurrence of Omicron mutation pairs at these sites simply reflects the rarity of these mutations within both assembled SARS-CoV-2 genome sequences and raw intrapatient sequence datasets (Figure 4).

**Figure 7.**
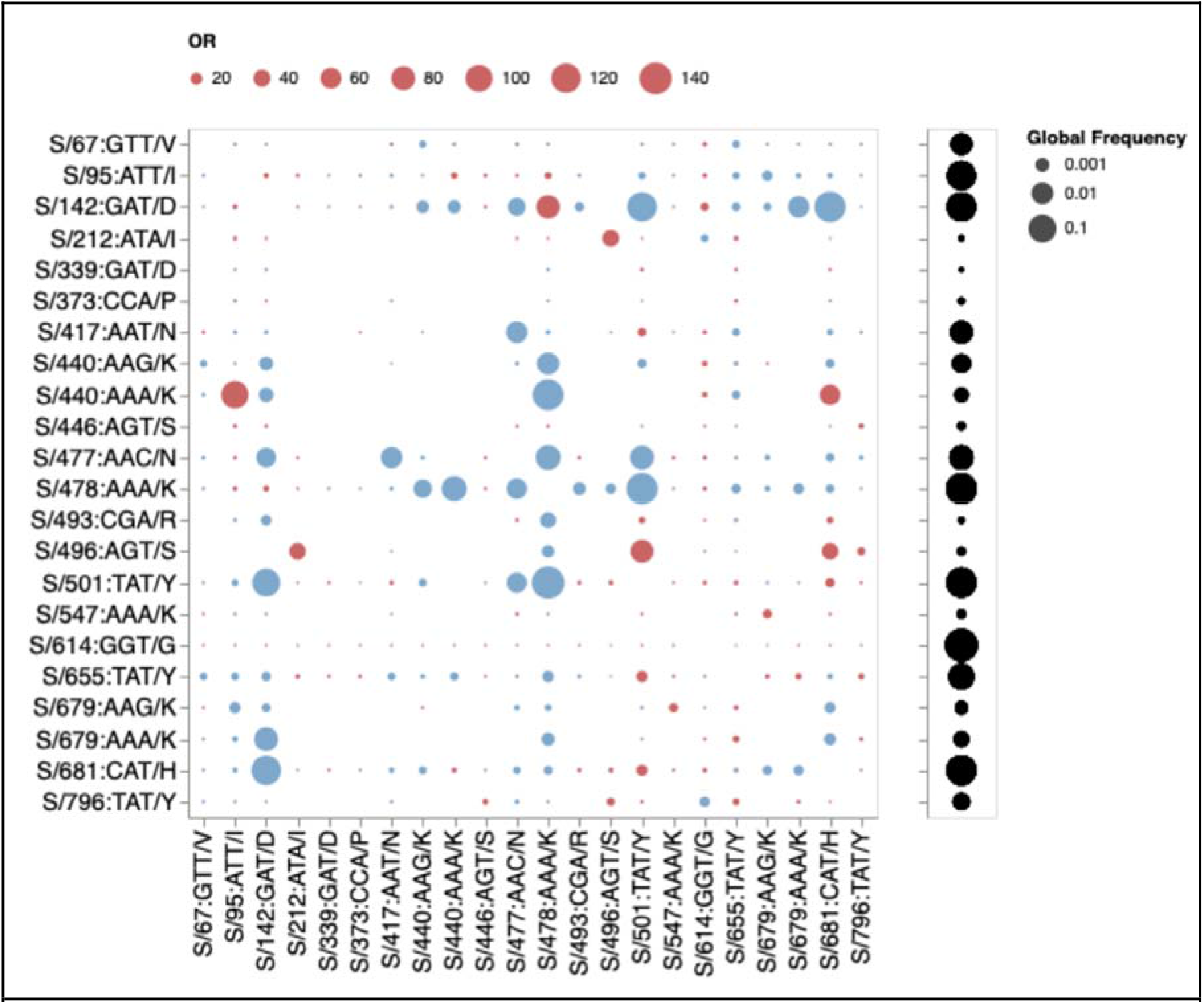
Patterns of co-occurrence of BA.1 amino-acid residues in circulating SARS-CoV-2 S-gene haplotypes from other lineages (data up to October 15, 2021). Only mutations occurring in at least 10 haplotypes are shown. All sequences having exactly the same S-gene sequence count as a single unique haplotype; instead of counting raw sequence numbers, this approach focuses on the number of unique genetic backgrounds in which pairs of codons co-occur. Circles show odds ratios for finding the mutation on the X axis when the mutation on the Y axis is also present (vs when it is not present). Red circles depict OR > 1, while blue circles 1/OR for OR < 1. Black circles on the right show the fraction of globally sampled SARS-CoV-2 S-gene haplotypes which carry the corresponding mutation.

### Evidence that cluster 1, 2 and 3 sites may be coevolving with other genome sites during the ongoing diversification of BA.1

If pairs of the 13 mutations in the three cluster regions are epistatically interacting we would expect that these mutations might show evidence of coevolution during the ongoing diversification of the BA.1 lineage. We therefore tested the 135247 BA.1 annotated S-gene sequences that were available in GISAID ^12^ as of 05 Jan 2022 for evidence that any of the 630 site pairs with sufficient evolutionary signal (at least two non-synonymous substitutions along internal branches of a subsampled tree of genetically unique S-gene sequences) were coevolving using a Bayesian graphical model method ^41^.

We found six pairs of sites to be coevolving with posterior probability (PP) ≥ 0.9 (Figure 8). Two sites in Cluster 1 (S/371 and S/375) share substitutions along three internal tree branches (in all cases reversions to Wuhan-Hu-1 S residues at both sites) with the LF → SS reversion pair at these sites having a co-occurrence log-odds [LOD] of 6.5. In cluster 2, S/493 co-evolves with S/496 and S/498; in both cases substitutions along two internal branches are shared, and in both cases these substitutions are reversions to Wuhan-Hu-1 residues (RS → QG; LOD = 6.6 and RR → QQ, LOD = 6.4). One of the two branches involves the reversion of all three residues. Two sites in cluster 3, S/856 and S/954, are detectably coevolving in that they share a KH → NQ substitution pair along one internal tree branch ( LOD = 8.2).

**Figure 8.**
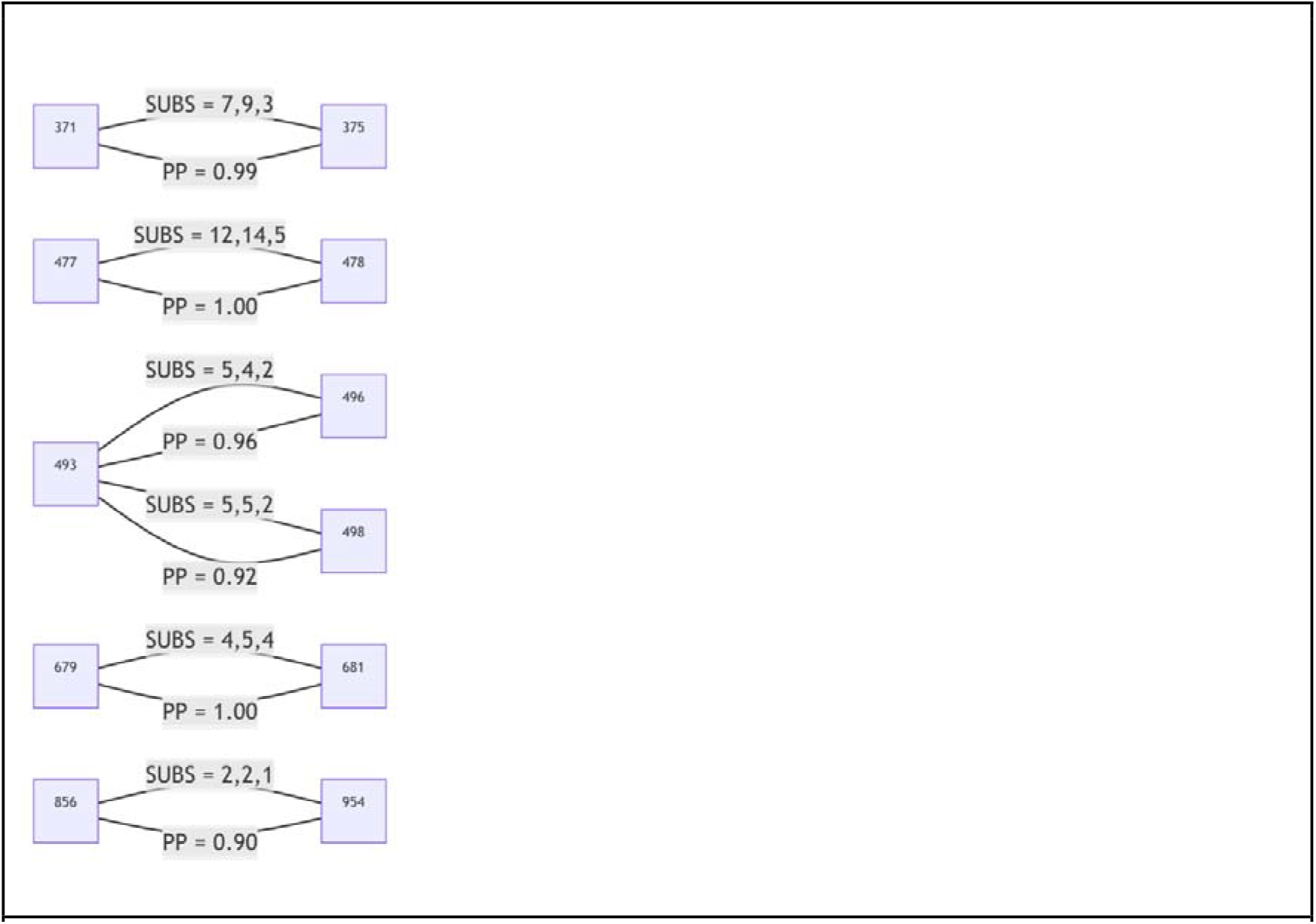
S-gene codon pairs that display substantial evidence of coevolution within the BA.1 lineage since the divergence of sampled BA.1 sequences from their most recent common ancestor. For SUBS = x,y,z: x = the number of non-synonymous substitutions likely occurring in the left codon along internal tree branches (i.e. where the mutant yielded multiple sampled and sequenced descendants): y = the number of non-synonymous substitutions likely occurring in the right codon along internal tree branches; and z = the number of non-synonymous substitutions likely occurring in both codons along the same internal tree branches. PP = posterior probability of conditional non-independence of substitutions at the two sites.

The detected coevolution between these site pairs supports the hypothesis that at least some mutations within each of the three cluster regions are epistatically interacting with one another, and, therefore, that the combined fitness impacts of the mutations in each of the cluster regions are likely more positive than the sum of the individual impacts of each mutation alone.

### How might mutations in the three cluster regions impact spike function?

Whether or not epistasis is restricted to a few site-pairs within the three cluster regions or is extensively operating between mutations within and/or between these regions, the amino acid changes caused by these mutations likely represent a substantial remodelling of two functionally important components of the BA.1 Spike: the receptor binding domain and the fusion domain.

The cluster region 3 encoded amino acid changes in the part of Spike that is responsible for membrane fusion suggest that the membrane fusion machinery of the BA.1 Spike may have been overhauled. These modifications possibly contribute to reduced TMPRSS2 mediated cleavage relative to Delta of BA.1 Spike at the polybasic S1/S2 cleavage site ^8^, reduced sensitivity to endosomal restriction factors (such as IFITM proteins) ^7^, and a shift in the preferred route of cellular entry from surface to endosomal ^6–8^: functionally important changes collectively resulting in a reduction relative to other SARS-CoV-2 lineages in the reliance of BA.1 on TMPRSS2 for cellular entry, a broadened cellular tropism, and a reduced propensity for infected cells to form syncytia ^7,8^.

The mutations in cluster regions 1 and 2 fall within the receptor binding domain (RBD) encoding part of the S-gene. These mutations, together with those at S/417, S/440, S/446 underlie an extensive remodelling of the ACE2 receptor binding surface ^10,30^; accommodating major changes in the way that Spike interacts with the ACE2 of humans and other animals ^7,9^.

Of the cluster 2 sites, all of which fall within the receptor binding motif encoding part of the RBD, only S/498 and S/505 show signs of the Wuhan-Hu-1 encoded amino acid state having been selectively favoured in the past (S/498 in SARS-CoV-2 and S/505 in nCoV). No signs of any positive selection at the other cluster 2 sites in SARS-CoV-2 implies that changes at these and the negatively selected sites in cluster 2 have likely not individually contributed to effective immune evasion since the start of the pandemic. Deep mutational scans (Figure 9; ^27^) have found little evidence that individual substitutions at S/505 have antigenic effects; S/496R and S/498R have only moderate antigenic effects, similar to those of the 501Y mutation. The exception that proves the rule that sites in this region might not be free to change in response to immune pressures is 493R. Given that 493R has a strong antigenic effect, if it was not under selective constraints to sustain optimal degrees of ACE2 interaction ^34^ it should (but does not) display at least intermittently detectable signs of positive selection.

**Figure 9.**
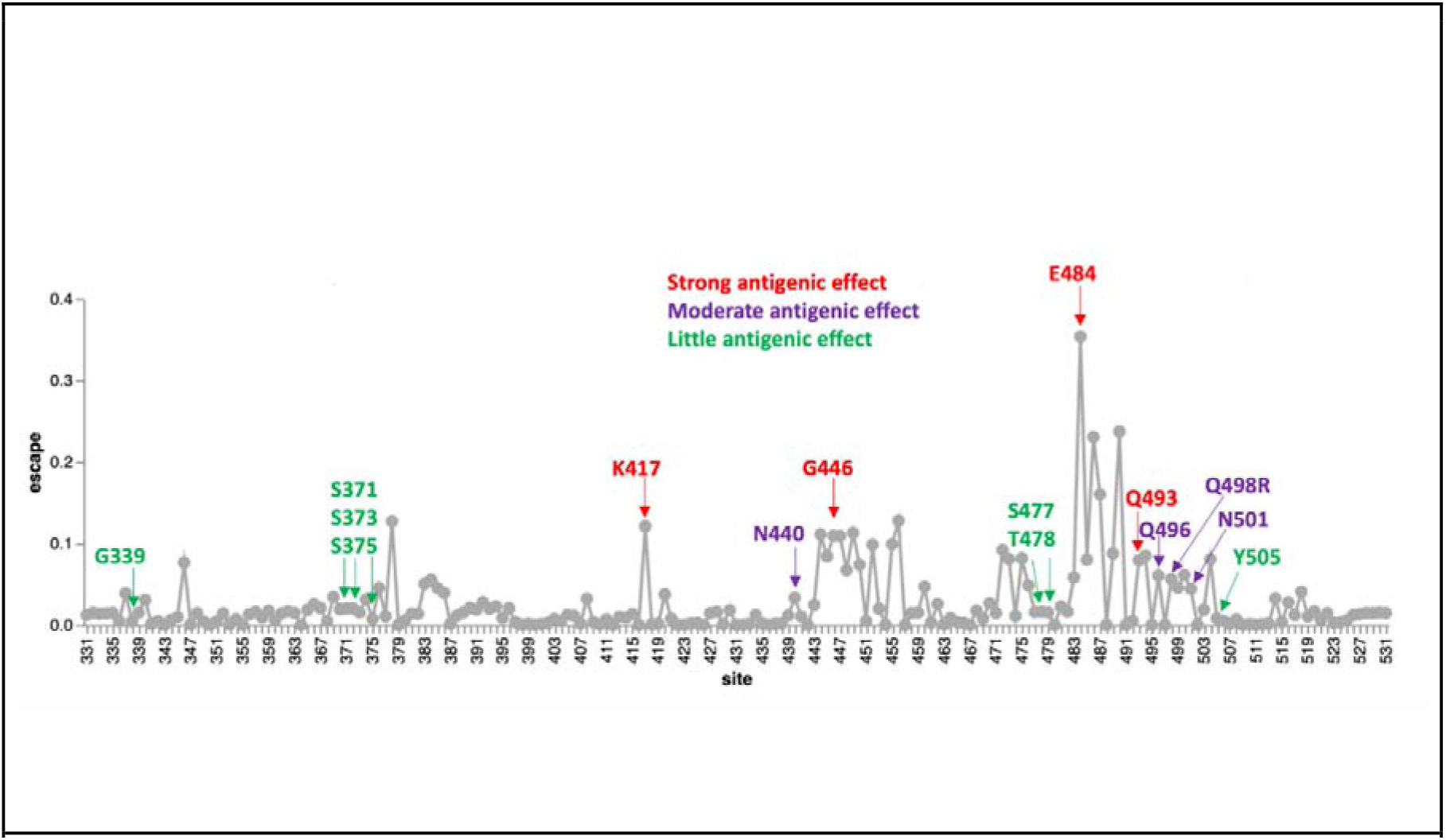
Experimentally measured effects of RBD mutations on binding of monoclonal antibodies at sites that differ between the BA.1 lineage viruses and Wuhan-Hu-1. The line plot shows antibody binding escape measured by deep mutational scanning of the Wuhan-Hu-1 RBD ^16^, averaged across 36 monoclonal antibodies (8 class 1, 13 class 2, 7 class 3, and 8 class 4 antibodies). Sites that are mutated in the BA.1 relative to Wuhan-Hu-1 are indicated and colored according to the predicted antigenic effect of mutations at that site (strong, moderate, or minimal). An interactive version of this plot is available at https://jbloomlab.github.io/SARS2_RBD_Ab_escape_maps/.

### How and why have so many apparently maladaptive mutations been assembled within Omicron?

Given the manifest viability of BA.1 and the other Omicron sub-lineages there is a pressing need to understand how and why they accumulated so many mutations that, on their own at least, are apparently either selectively neutral or maladaptive. The genetic distance between the Omicron sublineages and their nearest known SARS-CoV-2 relatives implies that the Omicron progenitor accumulated its unprecedented number of mutations during an extensive period of undetected replication. When accurate molecular clock estimates are obtained of both the time when Omicron last shared a common ancestor with other SARS-CoV-2 lineages, and the time when all the detected Omicron sublineages last shared a common ancestor, we will have upper and lower bounds on the amount of time it took for Omicron to assemble its complement of mutations.

The Omicron progenitor could have spent this period of intensive or prolonged evolution in a region that carries out minimal genomic surveillance and/or where access to, or utilization of, health care resources is low (the surveillance failure hypothesis). Alternatively, this viral evolution could have taken place within a long-term infection (or possibly serial long-term infections; the chronic infection hypothesis), or during spread within a non-human host population (the reverse-zoonosis hypothesis). Combinations of these evolutionary modes are also a possibility. We will only be able to distinguish between these hypotheses with more data.

Currently, the simple existence of three distinct Omicron lineages best supports the surveillance failure hypothesis at least for the latter stages of Omicron evolution following the divergence of the BA.1, BA.2 and BA.3 lineages from their most recent common ancestor. However, if similarly divergent SARS-CoV-2 variants are discovered in either longterm human infections or in other animal species, these would support the other hypotheses.

Relative to evolution during normal SARS-CoV-2 person-to-person transmission, evolution within the context of either long-term infections or an alternative animal host could potentially have occurred at an accelerated pace ^22,42^. In the context of either chronic infections of immunosuppressed individuals ^4,22,29^, or animals that naturally sustain long-term SARS-CoV-2 infections (such as may be the case for white tailed deer given the extraordinarily high frequencies of ongoing SARS-CoV-2 infections discovered in these ^43,44^), purifying selection may have been relaxed somewhat relative to that occurring during normal human-to-human transmission: enough so for genomes carrying suboptimal combinations of epistatically interacting mutations to remain viable while fitter combinations were discovered via additional mutations and genetic recombination. In addition, chronic infections are not impacted by the tight transmission bottlenecks that can stochastically purge nascent adaptive mutations during normal transmission ^45,46^.

Sequential cycles of immune surveillance and viral immune escape within a long-term infection could also potentially explain the mutation clusters without the need to invoke compensatory epistatic interactions between mutations. Specifically, the clustered mutation patterns in the Spike proteins of BA.1 and other Omicron sub-lineages are reminiscent of those seen in the HIV envelope protein as a consequence of sequentially acquired virus mutations that evade the progressively broadening neutralization potential of a maturing antibody lineage ^47^. While signs of negative selection at 9/13 of the mutated codons in the three cluster regions of Omicon are not entirely consistent with this hypothesis, the overwhelming contributor to these negative selection signals are the selective processes operating during normal short-term SARS-CoV-2 infections where the antibody-pathogen dynamics simply don’t have time to develop. It is possible that if purifying selection is relaxed at these sites during unusually prolonged infections, then neutralizing antibody evasion mutations might be tolerated. Even if purifying selection were not relaxed, however, during a chronic infection the potential long-term fitness costs that are incurred by highly effective immune evasion mutations might frequently be offset by the immediate fitness benefits of evading neutralization.

### It remains unclear whether mutations in cluster regions 1, 2 and 3 are showing signs of reversion

Whatever the process that yielded the three clusters of rarely seen mutations in the Omicron progenitor, now that it is being transmitted among people, any deleterious immune evasion mutations it has accumulated might be substantially less tolerable. Likewise, some of the mutations it may have accumulated during its adaptation to transmission in an alternative animal species would now also potentially be somewhat maladaptive. If the rarely-seen mutations at negatively selected sites in the RBD of BA.1 lineage viruses that are known to be targeted by neutralizing antibodies have begun reverting since BA.1 emerged, it would best support the chronic-infection hypothesis in that such reversions would imply a trade-off between intrahost replicative and/or movement fitness and immune evasion. Alternatively, if reversion mutations have occurred at BA.1 lineage virus receptor binding motif sites that are known to impact human ACE2 binding but which have minor antigenic impacts, this would better support the reverse zoonosis hypothesis.

Comparative evolutionary analyses focused on the BA.1 subclade of the SARS-CoV-2 phylogenetic tree revealed signatures of positive diversifying selection at 20 of the 28 S-gene codon sites that contain BA.1 lineage-defining mutations (Table 2, **bold**, deletions/insertions were not considered). Strong evidence of positive selection (FEL p < 0.001) was also detectable at several codon sites of the S gene that do not contain BA.1 lineage-defining mutations; most notably S/346 (R→K), S/452 (L→R) and S/701 (A→V). Amino acid changes encoded at all three of these codons are likely adaptive with S/R346K and S/L452R likely providing moderate degrees of escape from neutralizing antibodies ^16^, and S/A701V previously identified as one the 19 most adaptive Spike mutations within the context of N501Y carrying VOC lineages (Alpha, Beta and Gamma) ^15^.

**Table 2.**
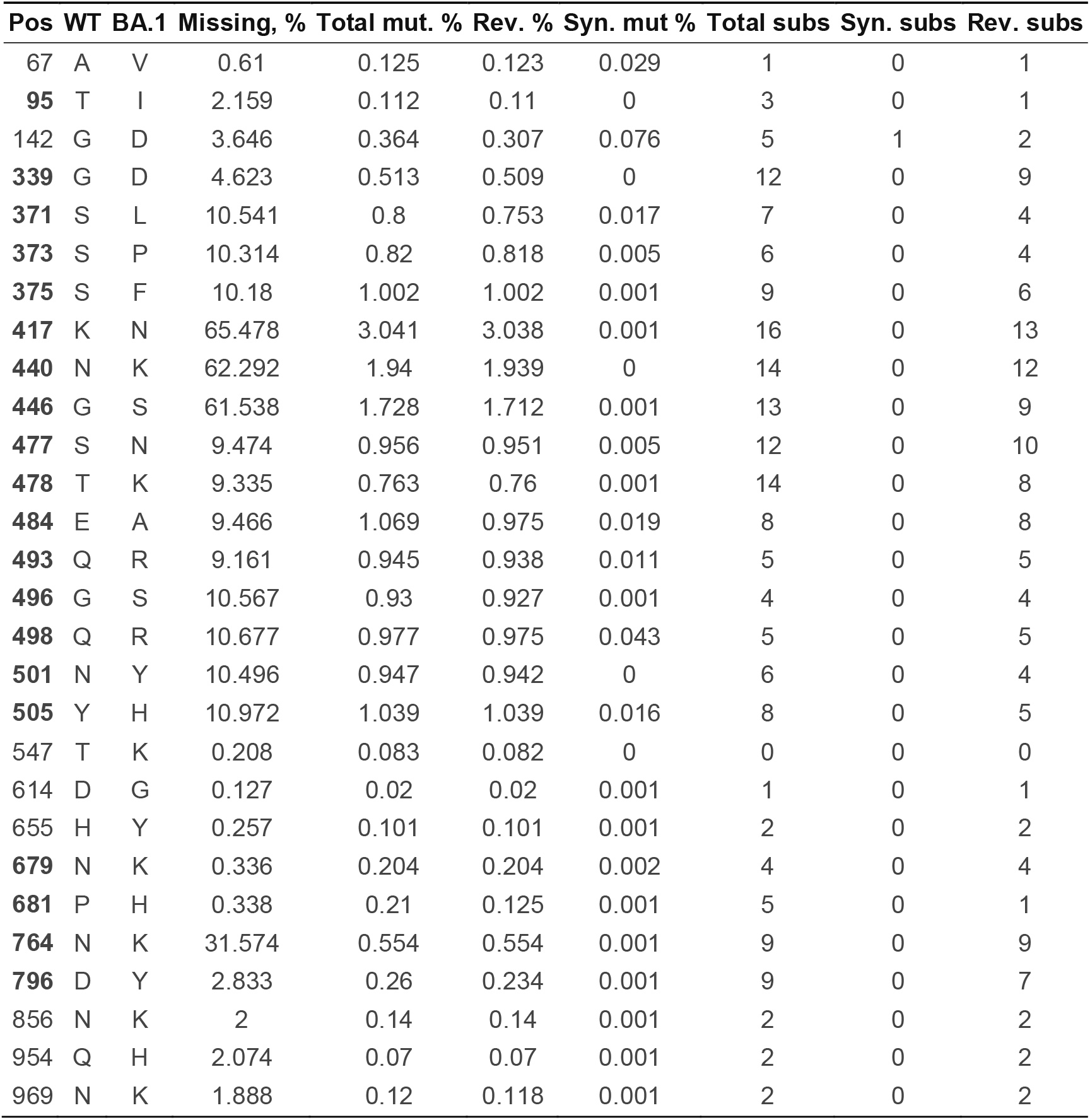
Evolutionary dynamics within BA.1 clade sequences at the positions of the S gene where BA.1 differs from the Wuhan-Hu-1 reference strain (WT) by an amino-acid change. **Missing, %** -- fraction of complete genomes in GISAID that have partially (e.g AAN) or completely (NNN) unresolved codons at this site. **Total mut, %** -- the fraction of sequences where there are mutations away from the BA.1 consensus codon (resolved codons only). **Rev, %** -- the fraction of sequences where there are mutations away from the BA.1 consensus back to the wildtype (WT). **Syn.mut, %** -- the fraction of sequences where there are synonymous mutations that maintain the BA.1 residue. **Total subs** -- the number of substitutions along internal branches of the BA.1 phylogeny which involve resolved nucleotides (based on the SLAC method); **Syn subs.** -- the number of substitutions that are synonymous for the BA.1 consensus residue; **Rev subs.** -- the number of substitutions that replace the BA.1 consensus residue with the WT residue. Bolded sites are those which are experiencing episodic positive selection along internal tree branches.

We found no molecular evidence for negative selection at any of these sites. At all sites, the vast majority of changes, measured either as fractions in all consensus genomes, or substitutions along internal branches of the phylogenetic tree of representative sequences, involve reversions to Wuhan-Hu-1 amino-acid states. At all sites, a fraction of sampled genomes have missing data (fully or partially unresolved nucleotides; Table 2). For key sites in RBD, this fraction is very high and, crucially, there is a strong correlation (R^2^ = 0.773) between the percentage missing data at a site and the number of reversion mutations inferred at that site (Figure 10). When multiplexing multiple samples in single sequencing runs, it is likely that known primer dropout issues for BA.1 sequences ^48^ can result in the amplification of environmental SARS-CoV-2 nucleic acid templates (e.g. from Delta lineages) that contaminate sample preparation laboratories and sequencing devices. When sequence reads derived from these contaminating templates are amplified to a similar degree to (or a greater degree than) BA.1 templates for a given region and are then used to assign nucleotide states in assembled genomes, apparent reversion mutations could result.

**Figure 10.**
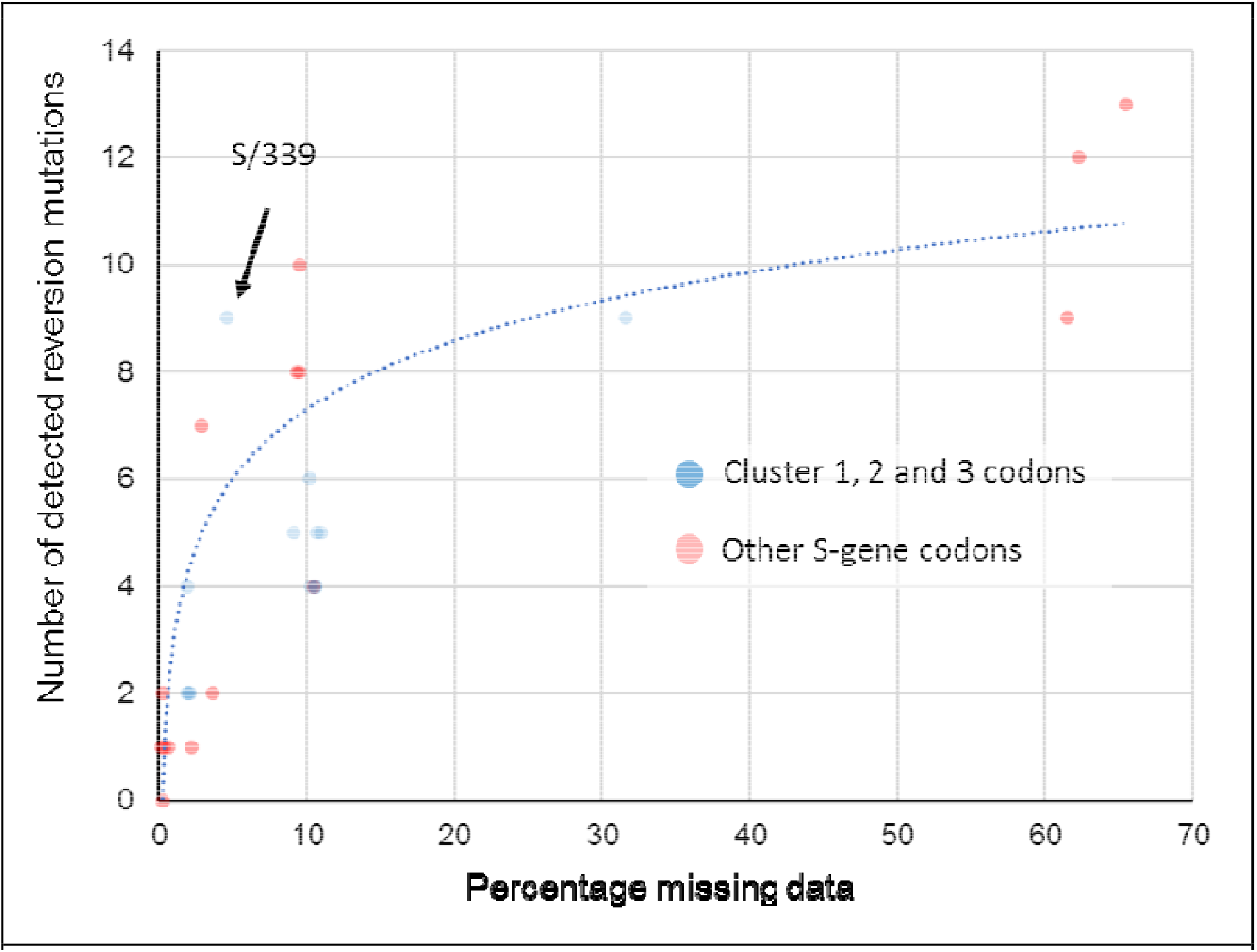
Association between the proportion of sequences with missing data at a BA.1 mutation site and the number of reversion mutations seen at that site. This significant association between missing data and reversion mutation counts (dotted blue trendline with Pearson’s R^2^ = 0.773; p < 0.01) is likely attributable to miscalled nucleotides at BA.1 mutation sites whenever read coverage is low during sequencing. Under conditions when PCR/sequencing primers are not optimal for the amplification of BA.1 sequence, non-BA.1 SARS-CoV-2 genetic material contaminating sequencing instruments and other laboratory equipment used for sample preparation, will occasionally yield more amplions/sequence reads than those from the intended BA.1 target sequences. Wherever the nucleotide states of these contaminant amplicons are different than those of the intended BA.1 target, they will frequently yield base miscalls during sequence assembly that, if the miscalled base corresponds with an ancestral state, will be misinterpreted as reversion mutations. Compared to BA.1 lineage-defining mutations in the S-gene at codon sites that are positively selected (red dots), the thirteen mutations at negatively selected or neutrally evolving cluster region 1,2 and 3 sites (blue dots) actually have a lower than average number of detectable reversion mutations (note how the blue dots predominantly fall below the blue trend line). Only one of these 13 mutations (at codon S/339) has a number of reversions that might be higher than expected given the percentage missing data for the codons where the mutations occur.

It is therefore unsurprising that for every BA.1 mutation in cluster regions 1, 2 and 3 we found multiple instances of reversions occurring along internal tree branches (a mean of 4.7 reversions per site; computed over internal tree branches in the reduced haplotype tree; Table 2). However, we noted that this pattern was also apparent for all of the other BA.1 spike mutations (a mean of 4.3 reversions per site): particularly so for the 15 BA.1 mutations falling within the RBD (mean of 5.3 for the cluster region 1 and 2 sites and 9.1 for the other sites). Further, of the 144 reversion mutations found across all of the S-gene, 13 (9.0%) were within clusters of three to four contiguous mutations: a degree of clustering that is significantly higher than would be expected for random independent mutations (permutation p-value < 0.001) (Figure 10). This pattern would, however, be expected with the widespread use of sequencing primers that are poorly suited to BA.1 sequencing.

When we account for the association between sequence coverage and reversion mutation counts, it is apparent that in the S-gene we do not see more reversion mutations at cluster region 1, 2 and 3 codon sites than at other BA.1 lineage-defining mutation sites (Figure 10). It therefore follows that, by this metric, the cluster 1, 2 and 3 mutations are, with the possible exception of that at S/339 (Figure 10), not obviously less adaptive during the ongoing diversification of BA.1 than are the other S-gene BA.1 lineage-defining mutations.

Despite not supporting one origin hypothesis over another, our inability to convincingly demonstrate unusually frequent reversions of cluster region 1, 2 and 3 mutations, remains consistent with the hypothesis that these mutations are broadly adaptive when they occur in the combinations found in BA.1 lineage viruses.

## Conclusion

Regardless of how the complement of mutations in the three cluster regions was assembled, their presence in BA.1 together with indirect evidence that the mutations are epistatically interacting is concerning. As with the concomitant emergence of the Alpha, Beta and Gamma VOCs in late 2020, part of the reason that the emergence of Omicron was a surprise is that the evolvability of SARS-CoV-2 is still deeply under-appreciated. It is becoming increasingly apparent that the evolutionary processes that yielded BA.1 involved balancing multiple fitness trade-offs: (1) between immune escape ^6,9,26,49,50^ and affinity for human and/or animal ACE2 proteins ^7–10,30^; (2) between efficient proteolytic priming with TMPRSS2 which expedites cellular entry via the cell surface ^6–8^ and increased resistance to endosomal restriction factors (such as IFITM proteins) which enable more efficient cellular entry via the endocytic route ^7^; (3) between preferred tropism for cells in the upper respiratory tract and preferred tropism for cells in the lower respiratory tract ^7,8^, and (4) between increased propensity for Spike monomers to switch from the down to up configurations and overall Spike trimer stability ^37,39^. Fortunately, the collection of mutations in BA.1 appear at present to have tilted the balance of these and other trade-offs towards the virus having decreased clinical severity in humans ^51,52^.

It remains unclear what roles epistatic interactions between the BA.1 S-gene cluster region 1, 2 and 3 mutations have played in resolving these trade-offs. It is evident, however, that the extensive mutational changes in BA.1 that have collectively yielded these resolutions are as similar to “normal” stepwise mutational changes seen in previous variants as antigenic shifts are to antigenic drifts ^53^. The evolutionary dynamics of the clustered rarely seen mutations in the RBD and fusion domains of BA.1 lineage viruses suggest that - rather than merely supporting minor tweaks in the antigenicity of Spike, its ACE2 binding affinity or its membrane fusion properties - these mutations are likely pivotal to the big observed shifts in how BA.1 Spike proteins function.

While a threat in its own right, BA.1 is also a warning. It demonstrates that complex evolutionary remodelling of important functional elements of SARS-CoV-2 are not just possible, but are potentially already occurring unnoticed in other poorly sampled lineages. We should not complacently assume that the balance of fitness trade-offs achieved by the extensively evolved VOCs that succeed BA.1 will be similarly tilted towards lower severity.

## Methods and Materials

### Global analyses of selection

Unless specified otherwise, all analyses were performed on single gene (e.g. S) or peptide products (e.g. nsp3); since genes/peptides are the targets of selection. Global SARS-CoV-2 gene/peptide datasets were compiled (from GISAID; ^12^), processed and analysed at monthly intervals for evidence of selection acting on individual codon sites as in ^15^. Results of these analyses at codons where Omicron mutations occur can be visualized using an Observable notebook at https://observablehq.com/@spond/sars-cov-2-selected-sites.

### Analyses of intrapatient SARS-CoV-2 diversity

Intrahost allelic variation seen at BA.1 amino acid mutation sites was analysed in 282788 annotated (i.e. with detailed associated metadata) publically available SARS-CoV-2 raw sequencing datasets from the UK, Greece, Estonia, Ireland and South Africa between March 2020 and September 2021 all of which were processed and analyzed using the standardized variant calling pipeline described in ^24^. All variant calling data for genomic sites where BA.1, 2, and 3 lineage defining mutations occur were extracted from processed datatsets available via ftp://xfer13.crg.eu/ and https://covid19.galaxyproject.org/genomics/global_platform/#processed-cog-uk-data and can be explored using the observable notebook at https://observablehq.com/@spond/intrahost-dashboard.

### Analyses of selection in sarbecoviruses related to SARS-CoV-2

The whole genome sequences of 167 members of the *Sarbecovirus* subgenus (including SARS-CoV and SARS-CoV-2 Wuhan-Hu-1; See https://docs.google.com/spreadsheets/d/1sSt7fRiBYeW9z5Amj1_OywHhfxCnZ2wqo9gnLKsq74c/edit?usp=sharing for the full list of accession numbers) were aligned using MAFFT (with the localpair option ^54^). GARD ^55^ was used on the whole-genome alignment to determine 26 recombination breakpoints based on which individual gene codon alignments were separated. Phylogenies for the resulting putatively non-recombinant codon alignments were reconstructed using IQTREE2 ^56^ (GTR+I+F+G4 model) and selection signals specific to the nCoV clade branches were inferred using the FEL ^13^ and MEME ^57^ methods as in ^58^. Results of these analyses for all gene regions can be explored using the observable notebook at https://observablehq.com/@spond/ncos-evolution-nov-2021.

### Analyses of selection in the BA.1 sublineage

Because the codon-based selection analyses that we performed gain no power from including identical sequences, and minimal power from including sequences that are essentially identical, we filtered BA.1 and reference (GISAID) sequences using pairwise genetic distances complete linkage clustering with the tn93-cluster tool (https://github.com/veg/tn93). All groups of sequences that were within D genetic distance (Tamura-Nei 93) of every other sequence in the group were represented by a single (randomly chosen) sequence in the group. We set D at 0.0001 for lineage-specific sequence sets, and at 0.0015 for GISAID reference (or “background”) sequence sets. We restricted the reference set of sequences to those sampled before Oct 15th, 2020.

We inferred a maximum likelihood tree from the combined sequence dataset using raxml-ng using default settings (GTR+G model, 20 starting trees). We partitioned internal branches in the resulting tree into two non-overlapping sets used for testing and annotated the Newick tree. Because of a lack of phylogenetic resolution in some of the segments/genes, not all analyses were possible for all segments/genes. In particular, this is true when lineage BA.1 sequences were not monophyletic in a specific region, and no internal branches could be labeled as belonging to the focal lineage.

We used HyPhy v2.5.34 (http://www.hyphy.org/) ^14^ to perform a series of selection analyses. Analyses in this setting need to account for a well-known feature of viral evolution ^41^ where terminal branches include “dead-end” (maladaptive or deleterious on the population level) ^14^ mutation events within individual hosts which have not been “seen” by natural selection, whereas internal branches must include at least one transmission event. However, because our tree is reduced to only include unique haplotypes, even leaf nodes could represent “transmission” events, if the same leaf haplotype was sampled more than once (and the vast majority were). The branches leading to these repeatedly sampled haplotypes were therefore also included in the analyses.

We performed an additional analysis on BA.1 sequences, which includes data available in GISAID up to Jan 5th, 2022. The workflow for intrahost gene analysis is as follows (code available at https://github.com/veg/omicron-selection; please note the scripts require the GISAID FASTA files and are not robust to changes in input format).

1. Obtain GISAID sequences annotated as BA.1
2. Map them to the reference S-gene using bealign (part of the BioExt Python package). `bealign -r CoV2-S input.fasta output. bam; bam2msa output.bam S.mapped.fasta`
3. Identify all sequences that are identical up to ambiguous nucleotides using tn93-cluster (these are the unique haplotypes). `tn93-cluster -f -t 0.0 S.mapped.fasta > S.clusters.0.json; python3 python/cluster-processor.py S.clusters.0.json > S.haplo.fasta`
4. Reduce the set of unique haplotypes to clusters of sequences that are all within 0.002 genetic distance of one another (`tn93-cluster -f -t 0.002 S.haplo.fasta > S.clusters.1.json; python3 python/cluster-processor.py S.clusters.1.json > S.uniq.fasta`
5. Identify and remove all sequences that are 0.0075 subs/site away from the “main” clusters (outliers/low quality sequences which result in long tree branches, or are possibly misclassified)
6. For each remaining sequence cluster, build a majority consensus sequence using resolved nucleotides (assuming there’s at least 3). Remove clusters that comprise fewer than three sequences. Add reference sequences for BA.2 and BA.3 to add in tree rooting.
7. Building an ML phylogeny using raxml-ng. Annotate BA.1 internal branches.
8. Gene-level tests for selection on the internal branches of the BA.1 clades using BUSTED^57^ with synonymous rate variation enabled.
9. Codon site-level tests for episodic diversifying (MEME) ^57^ and pervasive positive or negative selection (FEL) ^13^ on the internal branches of the BA.1 clade.
10. Epistasis/co-evolution inference on substitutions along internal branches of the BA.1 clade using Bayesian Graphical models ^41^.
11. We combined all the results using a Python script and visualized results using several open source libraries in ObservableHQ (https://observablehq.com/@spond/ba1-selection).

## Acknowledgements

We gratefully acknowledge all of the authors from the originating laboratories responsible for obtaining the specimens and the submitting laboratories where genetic sequence data were generated and shared via the GISAID Initiative, on which this research is based. The Network for Genomic Surveillance in South Africa is supported by the Strategic Health Innovation Partnerships Unit of the South African Medical Research Council, with funds received from the South African Department of Science and Innovation. DPM, RW and CW are funded by Wellcome (222574/Z/21/Z). SLKP and AN are supported in part by R01 AI134384 (NIH/NIAID). OAM was supported by the Wellcome Trust [206369/Z/17/Z]. SL and DLR are funded by the MRC (MC_UU_12014/12). JDB’s work is supported in part by R01 AI141707. JDB is an Investigator of the Howard Hughes Medical Institute. WM and BG receive funding from the European Union’s Horizon 2020 and Horizon Europe research and innovation programmes under grant agreements No 871075 (ELIXIR-CONVERGE) and 101046203 (BY-COVID). PL acknowledges funding from the European Research Council under the European Union’s Horizon 2020 research and innovation programme (grant agreement no.~725422 - ReservoirDOCS), from the Wellcome Trust through project 206298/Z/17/Z, and from NIH grants R01 AI153044 and U19 AI135995. RJW receives support from the Francis Crick Institute which is funded by Wellcome (FC0010218), MRC (UK) (FC0010218) and Cancer Research UK (FC0010218). He also receives support from Wellcome (203135) and the Medical Research Council of South Africa. For the purpose of open access the author has applied a CC-BY public copyright licence to any author-accepted version of this manuscript.

